# The mutational landscape of SARS-CoV-2 variants diversifies T cell targets in an HLA supertype-dependent manner

**DOI:** 10.1101/2021.06.03.446959

**Authors:** David J. Hamelin, Dominique Fournelle, Jean-Christophe Grenier, Jana Schockaert, Kevin Kovalchik, Peter Kubiniok, Fatima Mostefai, Jérôme D. Duquette, Frederic Saab, Isabelle Sirois, Martin A. Smith, Sofie Pattijn, Hugo Soudeyns, Hélène Decaluwe, Julie Hussin, Etienne Caron

## Abstract

The rapid, global dispersion of SARS-CoV-2 since its initial identification in December 2019 has led to the emergence of a diverse range of variants. The initial concerns regarding the virus were quickly compounded with concerns relating to the impact of its mutated forms on viral infectivity, pathogenicity and immunogenicity. To address the latter, we seek to understand how the mutational landscape of SARS-CoV-2 has shaped HLA-restricted T cell immunity at the population level during the first year of the pandemic, before mass vaccination. We analyzed a total of 330,246 high quality SARS-CoV-2 genome assemblies sampled across 143 countries and all major continents. Strikingly, we found that specific mutational patterns in SARS-CoV-2 diversify T cell epitopes in an HLA supertype-dependent manner. In fact, we observed that proline residues are preferentially removed from the proteome of prevalent mutants, leading to a predicted global loss of SARS-CoV-2 T cell epitopes in individuals expressing HLA-B alleles of the B7 supertype family. In addition, we show that this predicted global loss of epitopes is largely driven by a dominant C-to-U mutation type at the RNA level. These results indicate that B7 supertype-associated epitopes, including the most immunodominant ones, were more likely to escape CD8+ T cell immunosurveillance during the first year of the pandemic. Together, our study lays the foundation to help understand how SARS-CoV-2 mutants shape the repertoire of T cell targets and T cell immunity across human populations. The proposed theoretical framework has implications in viral evolution, disease severity, vaccine resistance and herd immunity.

## INTRODUCTION

As of May 2021, the COVID-19 pandemic, caused by the novel Severe Acute Respiratory Syndrome Coronavirus 2 (SARS-CoV-2), has led to upwards 3.4 million deaths and 165 million confirmed cases worldwide (https://coronavirus.jhu.edu/map.html), making vaccine development and deployment an urgent necessity (Callaway, 2020). As a result of unprecedent efforts, vaccines have been developed and licensed within a 1-year timeframe and are currently being widely distributed for mass vaccination (Krammer, 2020).

A clear understanding of the natural protective immune response against SARS-CoV-2 is essential for the development of vaccines that can trigger lifelong immunologic memory to prevent COVID-19 (Sette and Crotty, 2021; Stephens and McElrath, 2020). Since the start of the pandemic, numerous studies have investigated the association between COVID-19 clinical outcomes and SARS-CoV-2 specific antibodies and T cell immunity (Altmann and Boyton, 2020; Bert et al., 2020; Braun et al., 2020; Grifoni et al., 2020a; Long et al., 2020a, 2020b; Meckiff et al., 2020; Moderbacher et al., 2020; Sekine et al., 2020; Weiskopf et al., 2020). Memory may be a concern for SARS-CoV-2 specific antibodies, as they were recently shown to be present in convalescent COVID-19 patients in a highly heterogenous manner (Dan et al., 2021) and, in some cases, observed to be undetectable just a few months post-infection (Seow et al., 2020). In contrast, an increasing number of studies point CD4+ and CD8+ T cells as key regulators of disease severity (Liao et al., 2020; Moderbacher et al., 2020; Schub et al., 2020; Weiskopf et al., 2020; Zhou et al., 2020). Studies of convalescent COVID-19 patients have also shown broad and strong CD4+ and CD8+ memory T cells induced by SARS-COV-2, suggesting that T cells may provide robust and long-term protection (Dan et al., 2021; Peng et al., 2020). Similar observations have been made for the most closely related human coronavirus, SARS-CoV, for which T cells have been detected 11 years (Ng et al., 2016) and 17 years (Bert et al., 2020) after the initial infection, whereas antibodies were noted to be undetectable after 2-3 years (Liu et al., 2006; Tang et al., 2011; Wu et al., 2007). Thus, vaccines designed to produce robust T cell responses are likely to be important for eliciting lifelong immunity against COVID-19 in the general population.

To investigate how T cells could contribute to long-term vaccine effectiveness, precise knowledge about SARS-CoV-2 T cell-specific epitopes is of paramount importance (Liu et al., 2020). To this end, bioinformatics tools were developed to predict T cell-specific epitopes during the early phase of the pandemic (Grifoni et al., 2020b). A comprehensive map of epitopes recognized by CD4+ and CD8+ T cell responses across the entire SARS-CoV-2 viral proteome was also recently reported (Tarke et al., 2020). Notably, the structural proteins Spike (S), Nucleocapsid (N) and Membrane (M) were shown to be rich sources of immunodominant HLA-associated epitopes, accounting for a large proportion of the total CD4+ and CD8+ T cell response in the context of a broad set of HLA alleles (Tarke et al., 2021). To date (May 2021), ∼700 HLA class I-restricted SARS-CoV-2-derived epitopes have been experimentally validated (https://www.mckayspcb.com/SARS2TcellEpitopes/) (Quadeer et al., 2020).

T cell epitopes that have been mapped across the entire SARS-CoV-2 viral proteome are reference peptides that are unmutated because they have been predicted from the sequence of the original SARS-CoV-2 that emerged from Wuhan, China (Grifoni et al., 2020b). However, analyses of unprecedented numbers of SARS-CoV-2 genome assemblies available from large-scale efforts have shown that SARS-CoV-2 is accumulating an array of mutations across the world, leading to the circulation and transmission of thousands of variants around the globe at various frequencies, and hence, contributing to the global genomic diversification of SARS-CoV-2 (Dorp et al., 2020a; Korber et al., 2020; Laamarti et al., 2020; Mercatelli and Giorgi, 2020; Mercatelli et al., 2020; Popa et al., 2020). In this regard, recent data indicate that most recurrent mutations appear to be evolutionary neutral with no evidence for increased transmissibility (Dorp et al., 2020a). Nonetheless, it is important to highlight that those neutral mutations are associated with a remarkably high proportion of cytidine-to-uridine (C-to-U) changes that were hypothesized to be induced by members of the APOBEC RNA-editing enzyme family (Dorp et al., 2020a; Giorgio et al., 2020; Klimczak et al., 2020; Kosuge et al., 2020; Li et al., 2020; Matyášek and Kovařík, 2020; Rice et al., 2020; Simmonds, 2020; Wang et al., 2020). Since shown for other viruses (Grant and Larijani, 2017; Monajemi et al., 2014), we reasoned that the putative action of such host enzymes during the first year of the pandemic could lead to the large-scale escape from immunodominant and protective SARS-CoV-2-specific T cell responses, thereby potentially compromising their effectiveness to control the virus at the population-scale.

In this study, we report a comprehensive study of the global genetic diversity of SARS-CoV-2 to expose the impact of mutation bias on epitope presentation and HLA-restricted T cell response within the first year of the pandemic, from December 2019 to December 2020. More specifically, we asked the following questions: 1) What are the impact of SARS-CoV-2 prevalent mutations detected across the global human population on the repertoire of validated SARS-CoV-2 T cell targets, with specific emphasis on CD8+ T cell epitopes? and 2) Are mutational patterns in the genomic and proteomic composition of SARS-CoV-2 indicative of disrupted (or enhanced) epitope presentation and T cell immunity in human populations? By answering these questions, we provide a theoretical framework to understand how SARS-CoV-2 mutants have shaped T cell immunity to evade effective T cell immune responses at the population level during the first year of the pandemic, i.e. without mass vaccination-induced immune pressure on viral evolution and adaptation.

## RESULTS

### The global diversity of SARS-CoV-2 genomes influences the repertoire of T cell targets

As of May 2021, nearly 1.7M complete SARS-CoV-2 genome assemblies are publicly available via the Global Initiative on Sharing All Influenza Data (GISAID) repository. In the context of this large-scale effort, we performed a global analysis of SARS-CoV-2 genomes to assess whether mutations that emerged during the first year of the pandemic could disrupt HLA binding of clinically relevant SARS-CoV2 CD8+ T cell epitopes. First, we identified missense mutations by aligning 330,246 high-quality consensus SARS-CoV-2 genomic sequences (GISAID; December 31^st^ 2020, prior to mass vaccination) to the reference sequence, Wuhan-1 SARS-CoV-2 genome (**Figure S1**). We found a total of 13,780 mutations identified in at least 4 SARS-CoV-2 genomes/individuals from GISAID, including 1,721 unique amino acid mutations in the S protein, with D614G as the most frequent one (94%) (Korber et al., 2020) (**Table S1** and **Table S2**). Next, we implemented a bioinformatics pipeline to assess the impact of these mutations on HLA binding for 620 unique SARS-CoV-2 HLA class I epitopes that were recently reported to trigger a CD8+ T cell response in acute or convalescent COVID-19 patients (Quadeer et al., 2020; Tarke et al., 2020) (see Methods). On average, we found that the predicted binding affinity of 181 of these SARS-CoV-2 epitopes (30%) for common HLA-I alleles was reduced by ∼100-fold (**Table S3** and **Figure S1**). It is also apparent that mutations negatively impacted the HLA binding affinity of 56 (31%) and 19 (10%) CD8+ T cell epitopes located in the immunodominant S and N proteins, respectively (**Figure 1A,B**). Notably, a gap in the N protein, composed of a serine-rich region, is associated with higher mutation rate and a marked lack of predicted T cell epitopes and response (**Figure 1B**). Epitopes located in the RBD vaccine locus were also impacted by mutations (**Figure 1C**).

**Figure 1.**
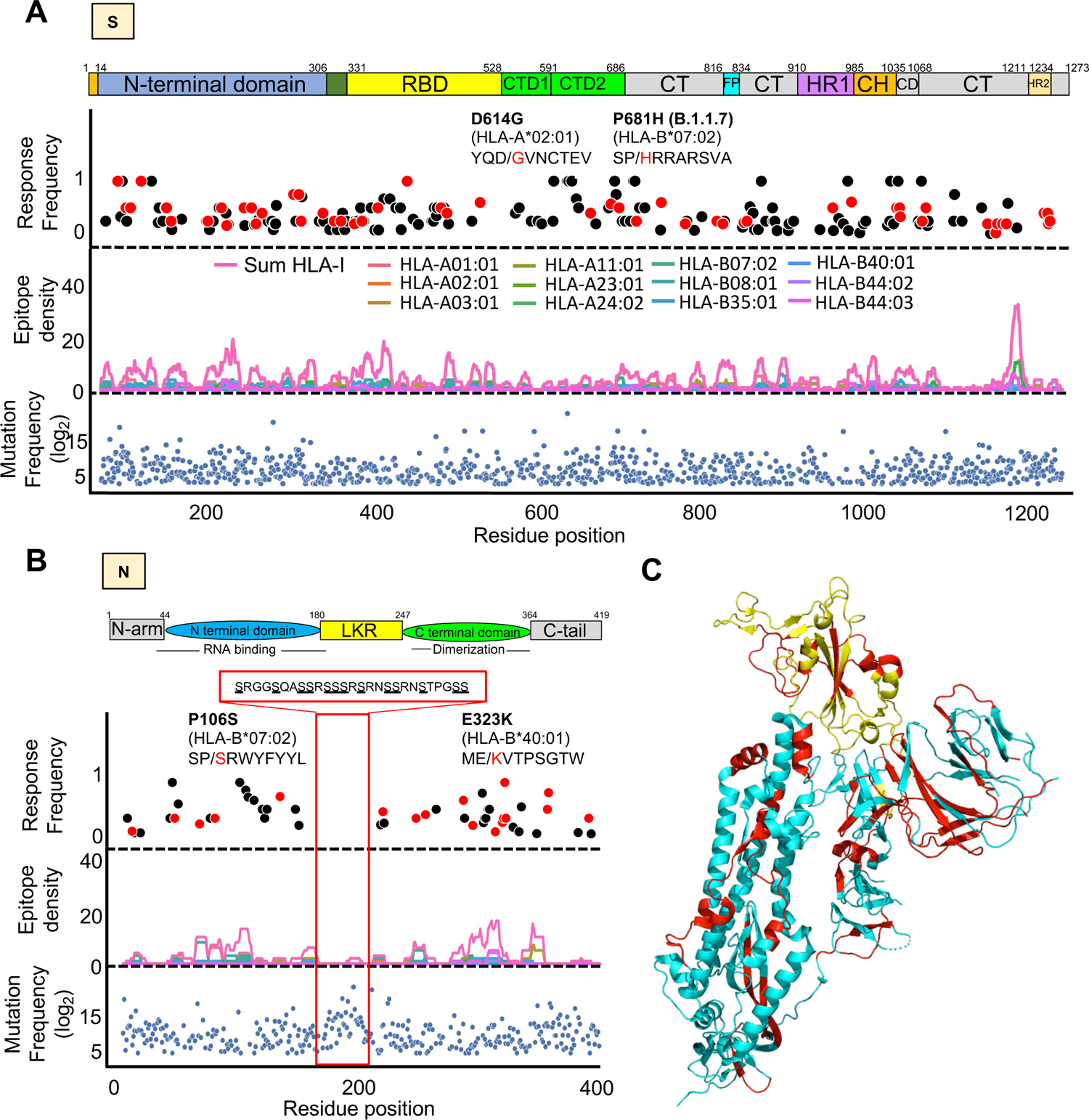
Distribution of CD8+ T cell epitopes and their mutated variants across the immunodominant S and N antigens. (**A, B**) Lower panel: blue dots showing all mutations that occurred in at least 4 SARS-CoV-2 genomes (GISAID). Middle panel: epitope density showing the overlap of HLA class I epitopes predicted within the 1st percentile for 12 queried HLA-I molecules. Upper panel: dots showing the frequency of CD8+ T cell response as determined from multiple studies aggregated in the database https://www.mckayspcb.com/SARS2TcellEpitopes as of January 2021. Red dots are mutated epitopes wherein the mutation event led to a predicted loss of binding. Sequences of specific epitopes are shown with the mutant amino acid in red. The red box in the N protein highlights a serine-rich region associated with no T cell response, low epitope density and high mutation frequency. (**C**) 3D structure of the Spike glycoprotein (Moderna Vaccine) and highlighted in yellow is the Receptor Binding Domain (Pfizer Vaccine). Shown in red are mutated epitopes wherein mutation events led to a predicted loss of HLA binding.

Loss of epitope binding for commonly expressed HLA class I molecules was validated *in vitro* for a subset of representative SARS-CoV-2 epitopes (**Figure S2**). Of relevance, we found that the common D614G mutation in the S protein is linked to a 15-fold decrease in the binding affinity for the mutated HLA-A*02:01 epitope YQ<u>G</u>VNCTEV when compared to the reference/unmutated epitope YQ<u>D</u>VNCTEV (**Figure S2A,B**). Interestingly, our analysis also identified a mutation in the HLA-B*07:02-restricted N105 epitope SPRWYFYYL, which is one of the most immunodominant SARS-CoV-2 epitope (Ferretti et al., 2020; Kared et al., 2021; Saini et al., 2021; Schulien et al., 2021; Sekine et al., 2020; Tarke et al., 2021). Although relatively rare (found in only two genomes), the mutation in the N105 epitope consists of PàS at anchor residue position P2 (P106S: S<u>P</u>RWYFYYL à S<u>S</u>RWYFYYL) (**Figure 1B**) and is predicted to decrease HLA epitope binding by 47-fold (**Figure 3D**), thereby likely reducing the breadth of the immune response in B*07:02 individuals carrying this mutation. Moreover, our global analysis validated the presence of two previously reported CD8+ T cell mutated epitopes (i.e. G<u>L</u>MWLSYFI à G<u>F</u>MWLSYFI, found in 38 genomes; and M<u>E</u>VTPSGTWL à M<u>K</u>VTPSGTWL, found in 23 genomes), which were shown to lose binding to HLA-A*02:01 and -B*40:01, respectively, in addition to disrupt epitope-specific CD8+ T cell response in COVID-19 patients (**Figure S3**) (Agerer et al., 2021). Together, these results demonstrate that mutations driving the global genomic diversity of SARS-CoV-2 can drastically disrupt HLA binding of clinically relevant CD8+ T cell epitopes, including epitopes encoded by the immunodominant S and N antigens, therefore affecting epitope-specific T cell responses in COVID-19 patients.

In addition to mutations leading to a loss of HLA epitope binding, we identified a significant number of mutations predicted to enhance the presentation of peptides by their respective HLA molecules, leading to a ‘Gain’ of binding (**Figure S4**). Because the unmutated epitopes are predicted to be non-HLA binders, these mutations were not searched against the list of known validated epitopes, which consist of strong-HLA binding reference epitopes. Whether SARS-CoV-2 mutations predicted to increase HLA epitope binding can enhance T cell responses to control the virus in COVID-19 patients remains to be determined experimentally.

### Amino acid mutational biases shape the global diversity of SARS-CoV-2 proteomes

While analysing the impact of the mutational landscape of SARS-CoV-2 on validated CD8+ T-cell epitopes, we observed that specific mutation types were over-represented while others were under-represented (**Figure S2C,D**). For instance, we found that 31% of the mutated epitopes were represented by a removal of proline residue (**Figure S2C,D**), leading to the hypothesis that such biases could originate from biases in the proteome of SARS-CoV-2 mutants. To further investigate whether specific amino acid mutational biases could be observed globally in the proteome of SARS-CoV-2 mutants, we asked whether certain amino acid residues were preferentially removed from, or introduced into the global proteomic diversity of SARS-CoV-2, thereby potentially diversifying CD8+ T cell epitopes in a systematic manner.

To test this, we computed all residue substitutions (amino acid removed and introduced) found in SARS-CoV-2 proteomes and calculated Global Residue Substitution Output (GRSO) values, i.e. the % difference in overall amino acid composition for individual amino acids (see Methods for details). GRSO values were computed for mutations found at various frequencies in GISAID (i.e. found in only 1 genome, 2 to 100 genomes, 100 to 1000 genomes and > 1000 genomes) (**Figure 2**). Interestingly, distinct mutational patterns at the amino acid level were observed amongst mutations detected in more than 100 genomes/individuals (**Figure 2**), referred in this study to as ‘prevalent mutations’ (see Methods and **Table S2**). Amongst those mutations, the amino acids alanine (A), proline (P) and threonine (T) were preferentially removed by 10.2% (p = 1.2×10^-13^), 9.1% (p = 1.6×10^-15^), and 10.5% (p = 1.3×10^-14^), respectively. In contrast, phenylalanine (F), isoleucine (I), leucine (L) and tyrosine (Y) were preferentially introduced by 13.4% (p = 2.0×10^-17^), 15.2% (p = 2.4×10^-17^), 4.3% (p = 6.3×10^-11^) and 5.0% (p = 7.0×10^-14^), respectively (**Figure 2**). Statistical significance of these GRSO values was assessed by generating simulated samples of 1000 SARS-CoV-2 genomes evolving under neutrality (N = 10 replicates) using the SANTA-SIM algorithm (Jariani et al., 2019) (see Methods for details). Of note, mutations that were detected in 2 to 100 individuals appeared significantly more neutral, with none of the mutational patterns enriched above the selected cut-off values (fold change > 4; p-value < 1×10^-11^). Thus, our results show that specific amino acid residues were preferentially removed or introduced in the proteome of SARS-CoV-2 mainly by prevalent mutations. Therefore, we introduce the notion that the global diversity of SARS-CoV-2 proteomes is shaped by specific amino acid mutational biases. Such biased amino acid composition generated by prevalent mutations may have a systematic impact on epitope processing and presentation to shape SARS-CoV-2 T cell immunity in human populations. To address this systematic impact, all downstream analyses described in this study were performed from the set of 1,933 prevalent mutations (>100 genomes) listed in **Table S2**.

**Figure 2.**
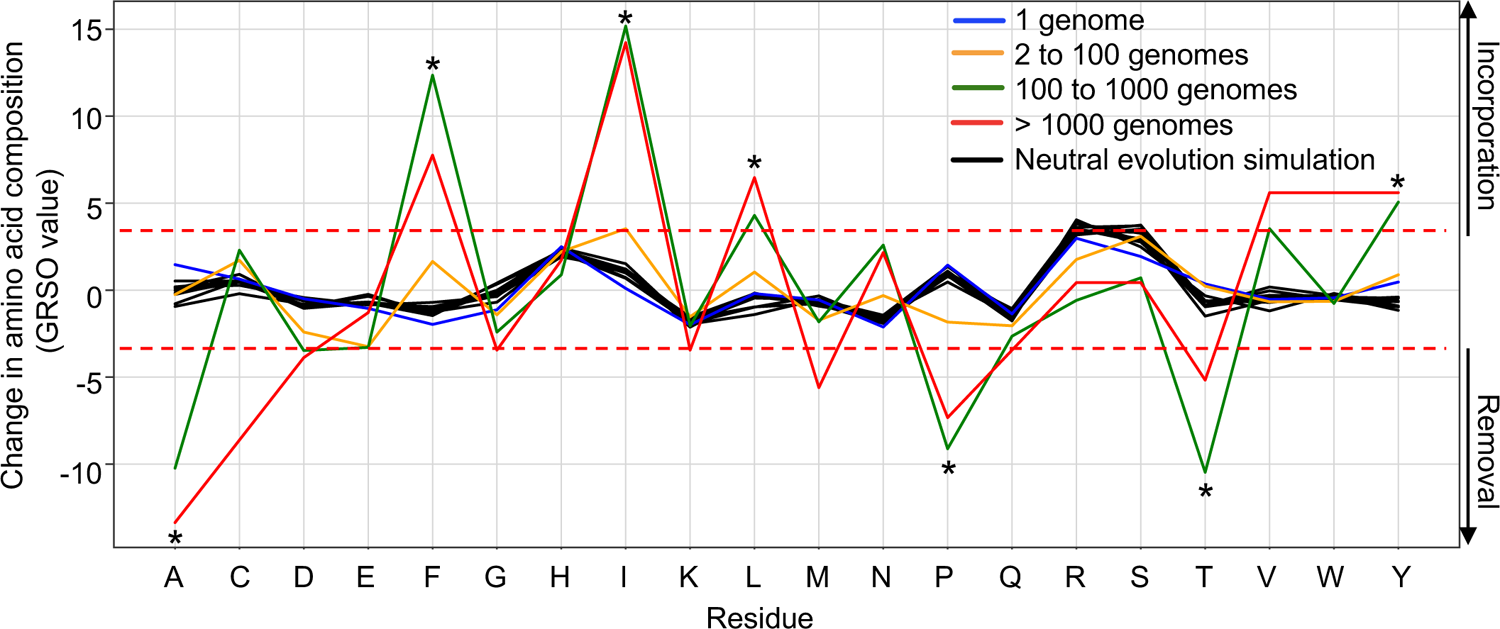
Global amino acid mutational biases in SARS-CoV-2 proteomes. A total of 330,246 SARS-CoV-2 genomes were translated into protein sequences and analyzed for the identification of any amino acid mutational bias. Amino acid residues (x-axis) that were removed and introduced in SARS-CoV-2 variants are presented by negative and positive %-difference in overall amino acid composition (GRSO values; y-axis), respectively. Analysis of mutational biases was performed for mutations occurring at various frequencies: 1 genome (blue line), 2 to 100 genomes (orange line), 100 to 1000 genomes (green line) and more than 1000 genomes (red line). Simulation of neutral evolution simulation (random mutations) were performed using the SANTA-SIM algorithm and serves as control for assessing the statistical significance of the observed pattern for individual amino acid residues. The dotted red lines show the cutoff values (fold change > 4; p-value < 1×10-11) that were used to define the residues that were preferentially removed or introduced (asterisk).

### Prominent removal of proline residues leads to a predicted global loss of epitopes presented by HLA-B7 supertype molecules

The association of peptides with the binding groove of HLA molecules largely relies on the presence of anchor residues, also known as peptide binding motifs (Falk et al., 1991). Hundreds of different peptide binding motifs have been reported over the last decades (Gfeller and Bassani-Sternberg, 2018). Overlapping binding motifs are qualified as “HLA supertypes” on the basis of their main anchor specificity (Greenbaum et al., 2011; Sidney et al., 2008). Of relevance here, proline acts as a critical anchor residue at position P2 for epitopes presented by HLA-B7 (B7) supertype molecules, which include a wide range of commonly expressed HLA-B alleles in humans, i.e. HLA-B*07, -B*15, -B*35, -B*42, -B*51, -B*53, -B*54, -B*55, -B*56, -B*67 and B*78 (Sidney et al., 2008). In fact, the B7 supertype covers ∼35% of the human population (Francisco et al., 2015). Hence, we reasoned that the global removal of proline residues observed in the proteome of prevalent SARS-CoV-2 mutants (**Figure 2**) could drastically compromise T cell epitope binding to B7 supertype molecules, thereby potentially interfering with SARS-CoV-2 T cell immunity in a relatively large proportion of the human population.

Due to the preferential removal of proline by prevalent mutations, we investigated the extent at which proline residues were substituted at anchor binding position P2 and, consequently, resulted in loss of epitopes presented by B7 supertype molecules. To answer this, we performed the following four steps: (i) We applied NetMHCpan 4.1 (Reynisson et al., 2020) using the reference and mutated SARS-CoV-2 genomes to generate a list of all possible reference/mutated peptide pairs (8-11 mers) predicted to bind 16 common HLA-B types that belong to the B7 supertype family (**Figure S5B**). (ii) We analyzed all reference/mutated peptide pairs, along with their differential predicted binding affinities to quantitatively identify HLA strong binder (SB) to non-binder (NB) transitions [(SB) NetMHCpan %rank < 0.5 to (NB) NetMHCpan %rank >2]. (iii) We categorized all peptide pairs based on the mutation type (amino acid X à amino acid Y) and the position of the mutation within the peptide sequence. (iv) Lastly, we quantified the number of reference/mutated peptide pairs and the associated fold-change in predicted binding affinity for each category. Our results show that prevalent mutations predicted to impact the presentation of peptides by the B7 supertype are dominated by PàL (p = 8.6×10^-35^) and PàS (p = 3.4×10^-24^) substitutions at anchor residue position P2 (**Figure 3A,B**). Reference/mutated peptide pairs from these categories were the most abundant, with > 250 mutated peptides per category (**Figure 3C**). PàL and PàS mutations resulted, on average, in a 61-fold reduction in predicted HLA binding affinity for a representative set of clinically validated CD8+ T cell epitopes (**Figure 3D**).

**Figure 3.**
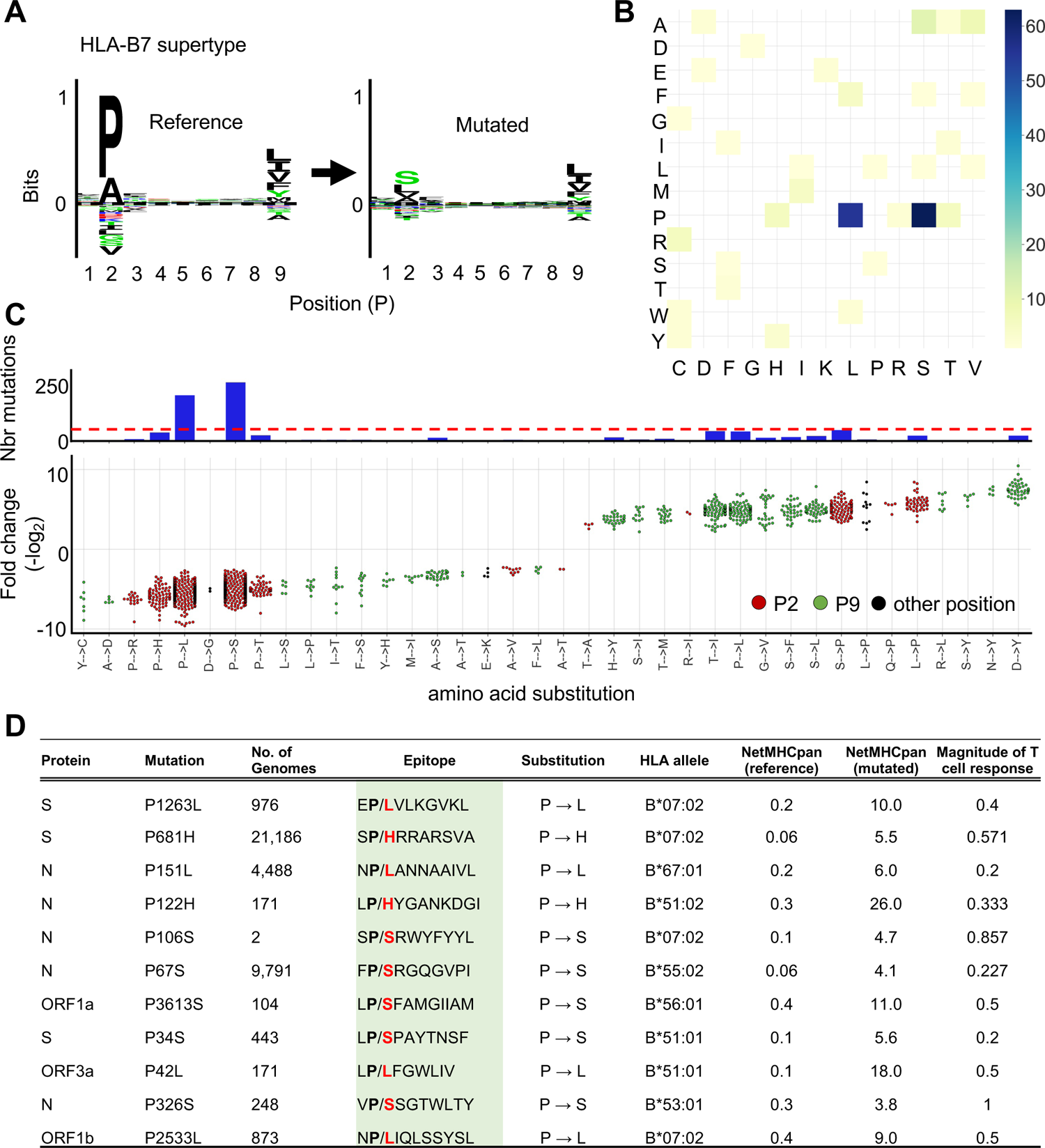
Mutation of proline at the anchor residue position for B7 supertype-associated epitopes. (**A**) (Left panel) Motif view of SARS-CoV-2 reference peptides predicted to bind B7 supertype molecules (HLA-B*07:02, -B*35:03, -B42:02, -B*5101, -B*53:01, -B*54:01, - B*55:01, -B*56:01, -B*67:01). (Right panel) Motif view of the corresponding mutated peptides. (**B**) Heat map showing the frequency of specific amino acid substitutions between reference and mutated peptides. (**C**) Graph showing the number of mutations (upper panel; y-axis) leading to specific amino acid substitutions (x-axis) at anchor residue positions P2 (red dots) and P9 (green dots) or elsewhere (black dots). Dotted red line indicate the cutoff used to define dominant substitutions. The lower panel shows fold changes for individual amino acid substitutions. (**D**) Representative examples of validated CD8+ T cell epitopes (https://www.mckayspcb.com/SARS2TcellEpitopes as of January 2021). Effect of the P→X substitutions on predicted epitope binding affinities (NetMHCpan 4.1 EL %Rank) are shown. T cell response data for reference epitopes extracted from https://www.mckayspcb.com/SARS2TcellEpitopes.

In addition to the dominant PàS/L substitution type, other PàX substitutions were observed. Interestingly, analysis of mutations found in the Pangolin B.1.1.7 variant (January 2021) showed that the P681H mutation found in the Spike protein led to disrupted association of the reference epitope S<u>P</u>RRARSVA for several HLA-B7 types. In fact, the P-to-H substitution resulted in a strong loss of epitope binding predicted for 7/16 HLA-B types tested. Thus, our results strongly suggest that biased substitutions of proline residues in the proteome of SARS-CoV-2 shapes the repertoire of epitopes presented by B7 supertype, including epitopes encoded by the genome of the B.1.1.7 variant. This finding let us to propose that mutation biases found in SARS-CoV-2 may contribute to CD8+ T cell epitope escape in a B7 supertype-dependent manner.

### The mutational landscape of SARS-CoV-2 enables disruption or enhancement of epitope presentation in an HLA supertype-dependent manner

We found that specific amino acid residues were preferentially removed (proline, alanine and threonine) or introduced (isoleucine, phenylalanine, leucine and tyrosine) in SARS-CoV-2 proteomes (**Figure 2**). Importantly, most of these amino acids act as key epitope anchor residues for multiple HLA class I supertypes (**Figure S5**). For instance, phenylalanine and tyrosine are key anchor residues for all known A*24 alleles of the A24 supertype family, whereas proline is known to play a critical role in the anchoring of epitopes to alleles of the B7 supertype family (**Figure 4**). Therefore, one would expect the introduction of phenylalanine and tyrosine in SARS-CoV-2 proteomes to facilitate peptide presentation by A24, whereas the removal of proline would disrupt peptide presentation by B7. With this concept in mind, we hypothesized that the distinct amino acid mutational biases found throughout prevalent SARS-CoV-2 mutations could systematically mold epitope presentation in an HLA supertype-dependent manner.

**Figure 4.**
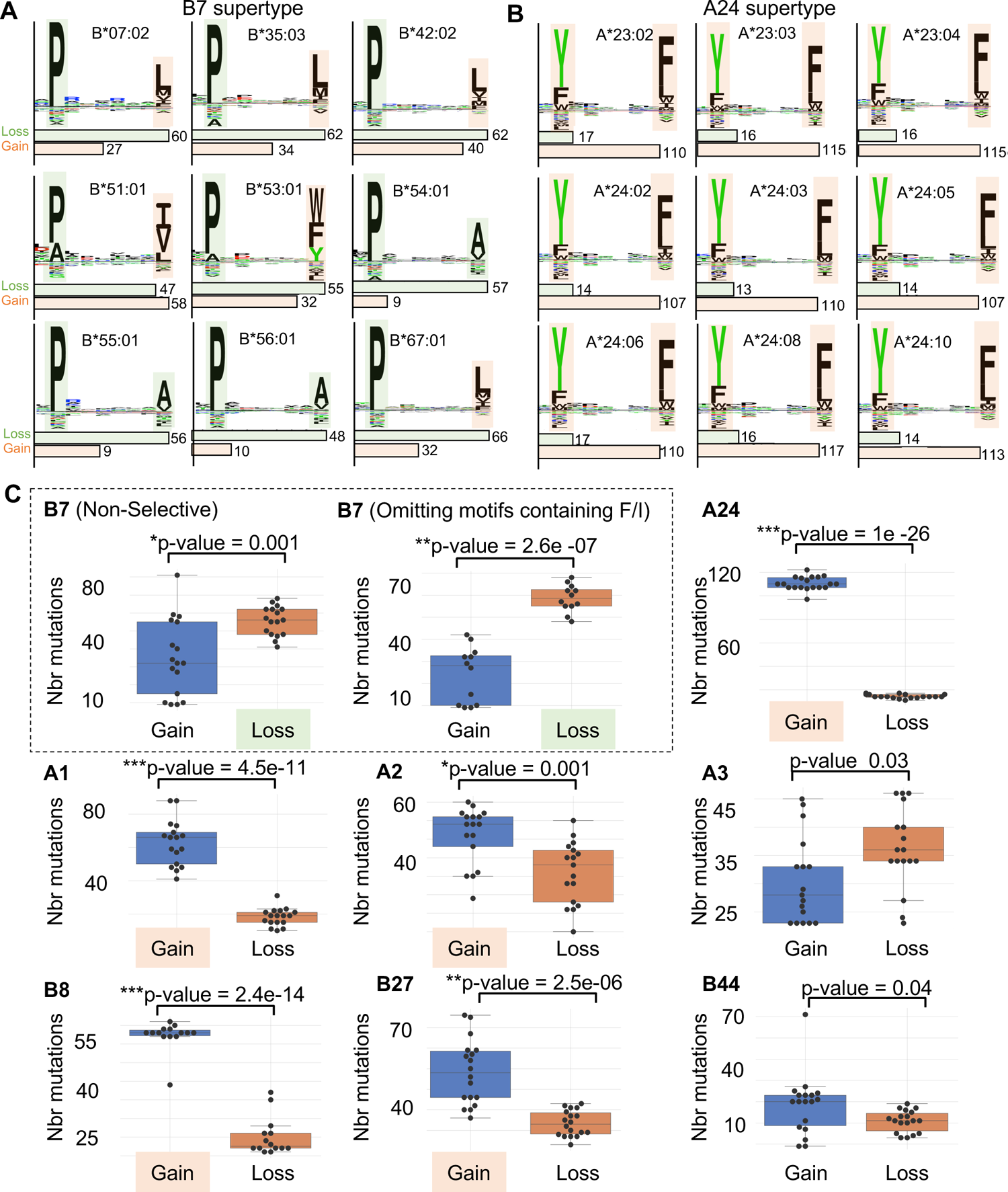
Loss or gain of SARS-CoV-2 mutated epitopes for different HLA class I supertypes. (**A, B**) Motif views showing established epitope binding motifs for different HLA-I alleles that belong to the HLA-B7 (A) and HLA-A24 (B) supertype family. Shaded squares highlight anchor residues that are preferentially removed (pale green) or introduced (pale orange) in SARS-CoV-2 proteomes (related to Figure 2), respectively. Histograms below the binding motifs indicate the number of frequent mutations (identified in at least 100 individuals) leading to the loss or gain of epitopes. (**C**) ‘Gain/Loss plots’ showing number of mutations (y-axis) leading to a preferentially loss (pale green) or gain (pale orange) of epitopes for different HLA class I supertypes. Each black dot represents the number of mutations associated with gain and loss of epitopes for a given HLA-I allele. Between 14 to 19 alleles per supertype (Figure S5) were used to generate the graphs and p-values (*p ≤ 0.001, **p < 1e-5, ***p < 1e-10).

In order to compare supertypes to each other, we generated a ‘Gain/Loss plot’ for each supertype assessed (**Figure 4C**). Gain/Loss plot were generated by computing the number of mutations that resulted in ‘Gain’ or ‘Loss’ of epitopes for representative class I alleles selected for each supertype (see methods for details). ‘Gain’ was assigned for mutated epitopes that were predicted to transit from non-HLA binders (NetMHCpan %rank > 2) to strong HLA binders (NetMHCpan %rank < 0.5), whereas ‘Loss’ was assigned for mutated epitopes that were predicted to transit from strong HLA binders to non-HLA binders. Surprisingly, our analysis shows that most supertypes preferentially gain new epitopes as a result of SARS-CoV-2 mutations: A1 (p = 4.5×10^-11^), A2 (p = 0.001), A24 (p = 1.0×10^-26^), B8 (p = 2.4×10^-14^), B27 (p = 2.5×10^-6^).

Interestingly, preferential loss of epitopes was only shown to be statistically significant for B7 supertype (p = 0.0012). Note that we explain the relatively low statistical value obtained for B7 supertype by the presence of isoleucine and phenylalanine (preferentially introduced in SARS-CoV-2 proteomes; see Figure 2) at anchor residue P9 for certain HLA types (namely HLAB*51:01 and HLA-B*53:01) (**Figure 4A**). In fact, omitting motifs containing isoleucine or phenylalanine increased the significance of epitope lost *versus* gained (p = 2.6×10^-7^) (**Figure 4C**). Together, our results show that the amino acid mutational biases that feature the global diversity of SARS-CoV-2 proteomes can positively or negatively affect binding affinities of mutated epitopes for a wide range of HLA class I molecules in a supertype-dependent manner.

### The C-to-U point mutation bias largely drives diversification of SARS-CoV-2 T cell epitopes

Next, we sought to better understand the genetic determinants that drive the association between epitope presentation and the amino acid mutational biases found in the SARS-CoV-2 population. To this end, we analyzed the abundance of all the possible nucleotide mutation types (i.e. A-to-C, A-to-G, A-to-U, C-to-A, C-to-G, C-to-U, etc.). This analysis indicates that C-to-U is the most common mutation type (43%), followed by G-to-U (28%), as well as A-to-G, G-to-A and U-to-C (from 9.7% to 11.6%) (**Figure S6A**), in line with observations made by others (Giorgio et al., 2020; Klimczak et al., 2020; Kosuge et al., 2020; Li et al., 2020; Matyášek and Kovařík, 2020; Rice et al., 2020; Simmonds, 2020; Wang et al., 2020).

Next, we aimed to determine the contribution of these different nucleic acid mutation types to the global mutational pattern observed at the amino acid level in Figure 2. To do so, we generated simulated population samples of 1000 SARS-CoV-2 genomes using SANTA-SIM (Jariani et al., 2019), applying various extents of mutational biases corresponding to the two most common mutation types observed (i.e. C-to-U and G-to-U). The resulting simulated viral populations were then analyzed to elucidate the global amino acid mutational pattern engendered by these simulated nucleic acid point mutation biases, and whether they recapitulate the observed patterns. Indeed, our data show that the mutational pattern resulting from the simulated C-to-U bias very closely mimicked the mutational pattern observed in the real-life dataset (**Figure 5A**). Namely, the *in silico* introduction of a C-to-U mutation bias resulted in the preferential removal of alanine, proline, and threonine, by 6.7% (p = 5.1×10^-11^), 6.9% (p = 1.2×10^-11^) and 8% (p = 4.8×10^-12^), respectively, as well as the introduction of isoleucine and phenylalanine by 8.2% (p = 1.3×10^-8^) and 5.2% (p = 4.3×10^-11^), respectively (**Figure 5A**). The G-to-U mutation bias also contributed to the introduction of isoleucine and phenylalanine (**Figure S6**). Together, these results show that the predominant C-to-U point mutations largely contribute to shaping the global proteomic diversity of SARS-CoV-2.

**Figure 5.**
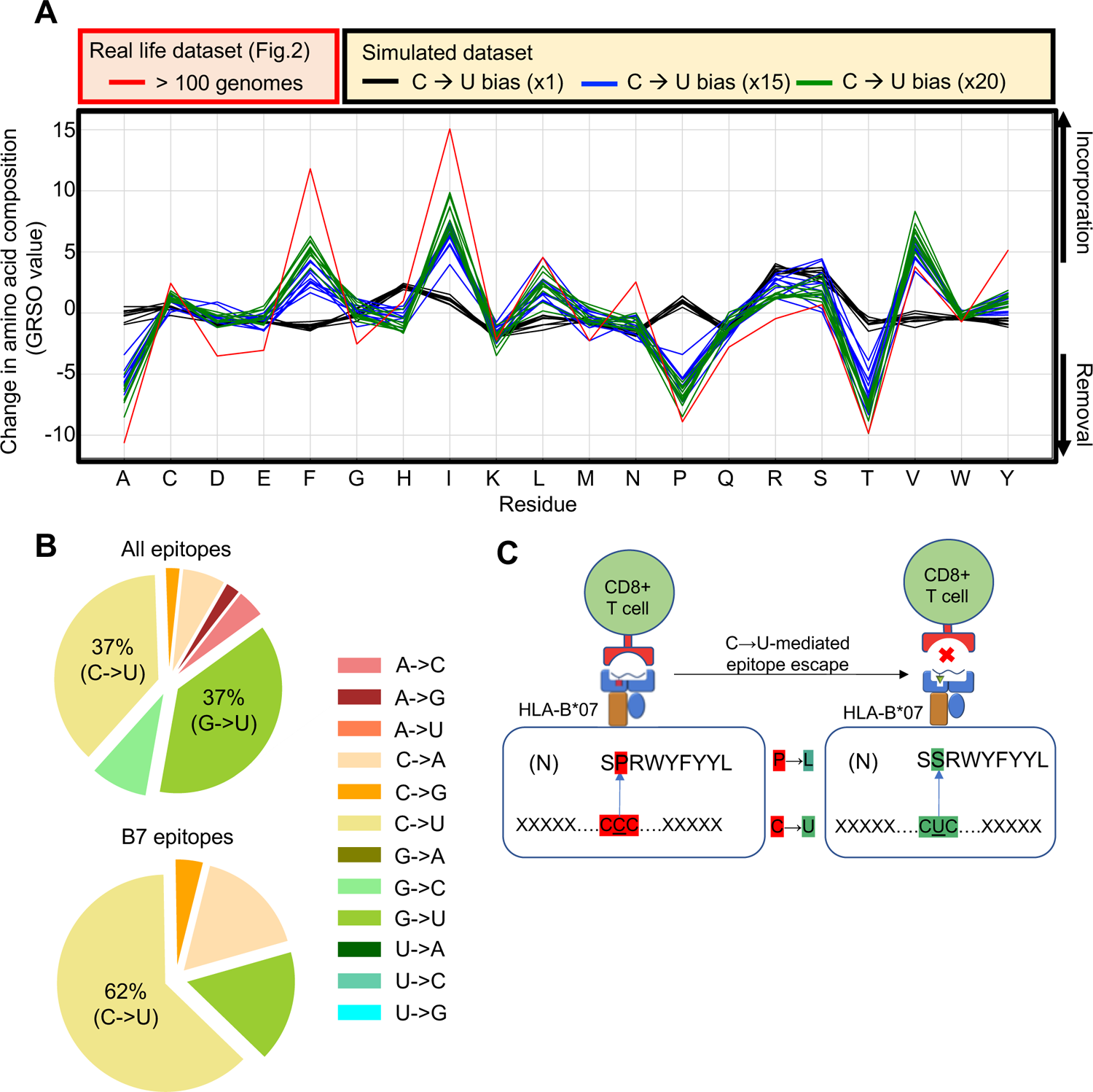
The C-to-U point mutation bias largely drives the diversity of SARS-CoV-2 proteomes and CD8+ T cell epitopes. (**A**) Comparison of global amino acid mutational patterns generated from real-life versus simulated SARS-COV-2 genomes. Amino acid residues (x-axis) that were removed and introduced in real-life versus simulated SARS-CoV-2 are presented by negative and positive %-difference in overall amino acid composition (GRSO values; y-axis), respectively. Evolution of SARS-CoV-2 was simulated by introducing various extents of C-to-U biases, i.e. x1, x15 and x20 (n = 10). The red line shows the pattern obtained from mutations identified in more than 100 SARS-CoV-2 genomes, related to Figure 2. (**B**) (Top) Pie chart showing the proportion of nucleotide substitution types from the list of validated CD8+ T cell epitopes in https://www.mckayspcb.com/SARS2TcellEpitopes as of January 2021. (Bottom) Pie chart showing the proportion of nucleotide substitution types from the list of validated CD8+ T cell epitopes specific to the B7 supertype. (**C**) Schematic illustrating the C-to-U-mediated epitope escape model. The observed mutation of the immunodominant SPRWYLFYYL epitope in the N protein is shown as an example.

Given the significant impact of the C-to-U point mutation bias on the amino acid content of SARS-CoV-2 proteomes, we reasoned that C-to-U could be the main driver shaping the repertoire and diversification of SARS-CoV-2 T cell targets in human populations, including targets presented by the particularly interesting B7 supertype molecules. To investigate this, we used all the SARS-CoV-2 CD8+ T cell epitopes that were experimentally validated using peripheral blood mononuclear cells (PBMC) of acute and convalescent COVID-19 patients (Quadeer et al., 2020; Tarke et al., 2020) and matched them with their corresponding nucleic acid sequence found in reference/mutated genome pairs. We then calculated the frequency of the various mutation types (i.e. A-to-C, A-to-G, A-to-U, C-to-A, C-to-G, C-to-U, etc.) coding for the mutated form of those clinically validated CD8+ T cell epitopes. Importantly, we found that C-to-U and G-to-U were the two main mutation types leading to mutated epitopes, both accounting for 37% of all mutation types amongst prevalent mutations (>100 individuals) (**Figure 5B**). Most strikingly, 62% of the prevalent mutations predicted to disrupt the presentation of epitopes by HLA alleles for the B7 supertype were found to derive from the C-to-U mutation type (**Figure 5B**). These results strongly suggest that the dominant C-to-U point mutation bias found amongst prevalent SARS-CoV-2 mutants has the potential to significantly contribute to shaping the repertoire of SARS-CoV-2 T cell epitopes in B7 supertype individuals across human populations.

Collectively, our study lets us to propose the model that C-to-U editing enzymes play a fundamental role in shaping the mutational landscape dynamics of SARS-CoV-2 CD8+ T cell targets in humans (**Figure 5C**), and hence, may contribute to molding T cell immunity against COVID-19 at the population level.

## DISCUSSION

Mutations contribute to the genetic diversity of SARS-CoV-2 and shape the progression of the COVID-19 pandemic (Dorp et al., 2020b, 2020a; Popa et al., 2020). T cells are key players controlling COVID-19 disease severity. Therefore, determining whether and how the mutational landscape of SARS-CoV-2 shapes or is shaped by HLA-restricted T cell response is fundamentally important. Traditionally, most studies have investigated how viral mutations are shaped by T cell response in the context of HLA-typed cohort patients. This type of approach sought to determine the evolutionary relationship between HLA genotypes and variants of long-standing viruses such as HIV-1 (Brumme et al., 2007; Kawashima et al., 2009) and influenza (Woolthuis et al., 2016). In the case of novel virus such as SARS-CoV-2, such a relationship remains to be established and does not constitute the scope of our work. Here, we rationalized that an alternative approach to interrogating SARS-CoV-2 epitope-associated variants is by investigating the global genomic and proteomic diversity of SARS-CoV-2 for any outstanding mutational biases, and then, assessing the relationship between such biases and epitope presentation for a broad set of HLA alleles. In other words, in this study, we did not seek to understand how viral mutations are shaped by T cell immunity, but rather to understand how mutational biases in SARS-CoV-2 may have shaped T cell immunity at the population level during the first year of the pandemic. This approach was possible thanks to an unprecedented number of SARS-CoV-2 genome sequences available for downstream analysis. Our approach is universal and could be applied to other epidemic or pandemic viruses in the future, given the development of distinct, prevalent mutational biases. Importantly, our global approach has led to several striking conclusions to help understand how the increasing genomic diversity of SARS-CoV-2 may shape T cell immunity in human populations. Our findings have important implications that are discussed below in the context of disease severity, viral evolution and vaccine resistance.

In this study, we found that prevalent SARS-CoV-2 mutations are governed by defined mutational patterns, with C-to-U being a predominant mutation type, as previously shown by others (Giorgio et al., 2020; Klimczak et al., 2020; Kosuge et al., 2020; Li et al., 2020; Matyášek and Kovařík, 2020; Rice et al., 2020; Simmonds, 2020; Wang et al., 2020). In fact, we show that the C-to-U mutation bias in SARS-CoV-2 genomes has a remarkably intimate relationship with the observed amino acid mutational biases, indicating that C-to-U mutations largely contribute to the global proteomic diversity of SARS-CoV-2. Most importantly, we show that this mutational bias leads to the preferential substitution of proline residues with leucine or serine residues in the P2 anchor position of SARS-CoV-2 CD8+ T cell epitopes, and hence, drastically compromise epitope binding to B7 supertype molecules, which represent ∼35% of the human population (Francisco et al., 2015). Therefore, the C-to-U mutational bias observed amongst prevalent mutants may partially disrupt SARS-CoV-2 T cell immunity in a very significant proportion of the human population. Noteworthy, this impact of C-to-U mutations on B7-depedent epitope escape was somehow predictable. In fact, proline residues originate from codons that are highly rich in C whereas serine and leucine residues originate from codons that are rich in both C and U. One could therefore predict, at least to some extent, that a strong C-to-U bias would lead to proline-to-leucine or proline-to-serine substitutions. Thus, this study highlights the impact of viral mutational biases and codon usage in shaping the diversity of CD8+ T cell targets. This being said, it is important to realize that we do not make the claim that the presence of proline-to-leucine or proline-to-serine mutations in the SARS-CoV-2 proteomes depend on patients being B7 supertype-positive, or that the B7 supertype drives the evolution of proline-to-leucine/serine mutations. We do, however, demonstrate that the prevalent mutations currently in circulation are enriched for proline-to-leucine/serine, and our *in silico* predictions suggest that the high occurrence of this mutation type leads to widespread hinderance of epitope presentation in B7 supertype-positive individuals.

A key question to address is to what extent does the C-to-U bias drives SARS-CoV-2 evolution and adaptation over the course of the ongoing pandemic. As proposed by others, the most likely explanation for the observed C-to-U bias is the action of the host-mediated RNA-editing APOBEC enzymes, a family of cytidine deaminases that catalyze deamination of cytidine to uridine in RNA (Dorp et al., 2020a; Giorgio et al., 2020; Kosuge et al., 2020; Olson et al., 2018; Salter et al., 2016). In this regard, APOBEC activity has been shown to broadly drive viral evolution and diversity, including in human immunodeficiency virus (HIV) (Albin et al., 2010; Cuevas et al., 2015; Haché et al., 2008; Jern et al., 2009; Peretti et al., 2018; Sadler et al., 2010; Wood et al., 2009). In fact, APOBEC-induced mutations driving the evolution and diversification of HIV-1 were shown to have an intimate relationship with T cell immunity (Kim et al., 2014; Wood et al., 2009). Notably, those studies have shown that the impact of APOBEC-induced mutations may result in either a decrease or increase of CD8+ T cell recognition, and that the direction of this response is dictated by the HLA context (Casartelli et al., 2010; Grant and Larijani, 2017; Kim et al., 2014; Monajemi et al., 2014; Squires et al., 2015; Wood et al., 2009). This is very much in line with our findings. Indeed, we showed that amino acid mutation biases in SARS-CoV-2 proteomes generally positively affect epitope binding for various HLA class I supertypes, and most strikingly for A24, whereas B7 is the only supertype negatively affected by the mutation biases given the markable loss of proline residues in SARSCoV-2 proteomes. Together, our results raise the important hypothesis that host-mediated RNA editing systems shape the repertoire of SARS-CoV-2 T cell epitopes in a positive and negative HLA-dependant manner.

Another question is whether populations of B7 supertype individuals represent an advantageous reservoir for the virus to evolve toward more transmissible variants. As the genetic diversity of the SARS-CoV-2 population continue to increase, and as new variants emerge, our global analysis suggests that the probability for SARS-CoV-2 epitopes to escape CD8+ T cell immunosurveillance is much higher in B7 individuals compared to A24 individuals. In fact, a slower T cell response dynamic to control SARS-CoV-2 infection in B7 individuals may offer a selective advantage for the virus to evolve. In this regard, we noted that the B.1.1.7 variant lost the B7 supertype-associated epitope S<u>P/H</u>RRARSVA as a result of a proline-to-histidine substitution.

While genomic surveillance is ongoing in different regions of the world, measuring the level of transmission of the B.1.1.7 variant within geographical regions of the world with low B7 population densities and high A24 population densities (in Asia) or the opposite trend (in Sub-Saharan Africa) (http://www.allelefrequencies.net/top10freqs.asp) may provide insights into this concern. As new variants of concern continue to emerge and as new epitope data are continuously being generated (Grifoni et al., 2021), another interesting avenue would be to study the mutational patterns of those emerging variants and assess whether and how the potential loss of B7-associated epitopes in those specific variants impact T cell response in infected patients. Understanding the impact of losing several subdominant B7-associated epitopes versus one single immunodominant epitope could also be investigated in the context of those variants. In this regard, a particular attention was allocated in our study to the B*07:02-restricted N105 epitope SPRWYFYYL. This epitope is of high interest as its immunodominance was experimentally demonstrated in many independent studies (Ferretti et al., 2020; Kared et al., 2021; Saini et al., 2021; Schulien et al., 2021; Sekine et al., 2020; Tarke et al., 2021). Precisely, we found a rare mutation consisting of PàS at P2 of this epitope (S<u>P</u>RWYFYYL à S<u>S</u>RWYFYYL). Its occurrence was predicted to result in the complete abrogation of binding of the epitope to B*07:02, thereby likely reducing the breadth of the immune response in individuals carrying this mutation. As such, we advise the community to carefully monitor this mutation in subsequent months. Moreover, it is also possible that B7 individuals respond less efficiently to the currently available vaccines, as genetic variants promoting B7 escape might favorably emerge in the future. The B7 supertype could therefore potentially represent a biomarker of vaccine resistance.

In summary, our study shows that mutation biases in the SARS-CoV-2 population diversify the repertoire of SARS-CoV-2 T cell targets in humans in an HLA-supertype dependent manner. Hence, we provide a foundation model to help understand how SARS-CoV-2 may continue to mutate over time to shape T cell immunity at a global population scale. The proposed process will likely continue to influence the evolution and diversification of SARS-CoV-2 lineages as the virus is under tremendous pressure to adapt in response to mass vaccination.

### LIMITATIONS AND FUTURE DIRECTIONS

Our analyses focused on class I molecules for which predictors are established to be more accurate in comparison with class II. HLA-C and non-classical HLA were not included in this study. Predictions were performed on the most common HLA class I alleles and rare HLA alleles were not included. Study has been performed using the GISAID dataset available in December 31^st^ 2020, i.e. first year of the pandemic, before mass vaccination. Our epitope binding results rely on *in silico* predictions using a method that has been widely benchmarked, but is designed to predict peptide presentation rather than immunogenicity. Follow up experiments would need to be performed to further validate the proposed model. Priority follow up studies are 1) to investigate T cell response to SARS-CoV-2 mutants in large cohorts of B7 supertype-positive versus negative patients, and 2) to determine the direct role of APOBEC family proteins in modulation of SARS-CoV-2-specific T cell immunity. Moreover, this study lays the foundation to understand the evolutionary dynamics of pandemic viruses with a time 0 / no vaccine-induced immune pressure start point. Employing SARS-CoV-2 as model provides an opportunity in future studies to look at the dynamic of the relationship between mutational patterns and HLA-restricted T cell immunity in real-time. Kinetic analyses using the latest GISAID datasets, which now include 1.7M SARS-CoV-2 genomes as of May 2021, may lead to additional insights in this regard.

## ACKNOWLEDGMENTS

We thank Drs. Alessandro Sette, John Sidney and Alba Grifoni (La Jolla Institute for Immunology, USA) for helpful discussions. This study was supported by funding from the Fonds de recherche du Québec – Santé (FRQS), the Cole Foundation, CHU Sainte-Justine and the Charles-Bruneau Foundations, Canada Foundation for Innovation, IVADO COVID19 Rapid Response grant (CVD19-030), Montreal Heart Institute Foundation, the National Sciences and Engineering Research Council (NSERC) (#RGPIN-2020-05232) and the Canadian Institutes of Health Research (CIHR) (#174924). K.K. is a recipient of IVADO’s postdoctoral scholarship (#4879287150). D.F. is a BioTalent awardee. E.C. and J.H. are FRQS Junior 1 Research Scholars.

## AUTHOR CONTRIBUTIONS

Conceptualization: D.H., J.H., and E.C.; Data Curation and Bioinformatic Analysis: D.H., D.F., J-C.G., F.M, K.K., and P.K.; Formal Analysis: D.H., and D.F.; Investigation: D.H., D.F., J.S., J-C.G., K.K., J.D.D., F.S., P.K., I.S., H.D., S.P., J.H., and E.C.; Writing – Original Draft: D.H., and E.C.; Writing – Review & Editing: D.H., D.F., J.S., J.S., J-C.G., F.M., K.K., P.K., J.D.D., F.S., I.S., M.S., H.S., H.D., S.P., J.H., and E.C.; Supervision: J.H., and E.C.; Funding Acquisition: J.H., and E.C.

## STAR METHODS

### RESOURCE AVAILABILITY

#### Lead Contact

Further information and requests should be directed to the lead contact, Dr. Etienne Caron

### Materials Availability

This study did not generate new unique reagents.

### Data and Code Availability

All sequence data used here are available from The Initiative for Sharing All Influenza Data (GISAID), at https://gisaid.org/. The user agreement for GISAID does not permit redistribution of sequences, but researchers can register to get access to the dataset. Code to create the alignments, to predict mutated and unmutated HLA-I peptides, and to perform the global analysis of SARS-CoV-2 proteomes are available at https://github.com/CaronLab.

## METHOD DETAILS

### Identification of SARS-CoV-2 mutations

All SARS-CoV-2 nucleotide sequences were acquired from the GISAID on 31/12/2021. A total of 330,246 SARS-CoV-2 sequences spanning 143 countries were acquired and analyzed. All sequences isolated from animals (including viral RNA isolated from bat, pangolin, mink, cat and tiger) were removed from the list and only high-quality sequences were further analysed. Consensus sequences were aligned to the reference sequence, Wuhan-1 (NC_045512.2) using minimap2 2.17-r974. All mapped sequences were then merged back with all others in a single alignment bam file. The variant calling was done using bcftools mpileup v1.91 in a haploid calling mode. Sequences were processed by batches of 1000 to overcome technical issues with very low-frequency variants. With the variant calling obtained for each batch, vcf-merge (from the vcftools suite) was used to merge all the variant calls across the entire dataset. A total of 24,220 variants in at least two consensus sequences were identified. Mutations appearing in only one genome were excluded as they are likely enriched for sequencing errors. A list of all missense mutations considered in our analyses is provided in **Table S1**. The 1,933 prevalent mutations observed in more than 100 genomes are also clearly shown in **Table S2**.

### Prediction of mutated and reference CD8+ T-cell epitopes

Prediction of CD8+ T cell epitopes was carried out using netMHCpan 4.0 EL (Reynisson et al., 2020). For each unique missense mutation, short sequence windows consisting of 14 amino acids on either side of the mutation site were generated, containing either the reference or mutated amino acid. Working from the resulting 29-residue sequence windows (mutation +/- 14 residues), 811mers were predicted against the 12 most frequent HLA alleles within the global population (HLA-A*01:01, HLA-A*02:01, HLA-A*03:01, HLA-A*11:01, HLA-A*23:01, HLA-A*24:02, HLA-B*07:02, HLA-B*08:01, HLA-B*35:01, HLA-B*40:01, HLA-B*44:02, and HLAB*44:03). Briefly, the NetMHCpan 4.0 EL method relies on a neural network trained on both binding affinity as well as eluted ligand data to produce a likelihood score for a peptide to be an eluted ligand for the indicated HLA types. The likelihood score consists of a percentile rank (%rank) wherein predicted (weak) binders obtain a %rank below 2.0, whereas strong binder (SB) obtain a %rank below 0.5. Using this ranking system, only mutation-containing peptides where the mutated and/or the reference peptide were ranked as SB were considered for further analyses. Mutations causing percentile ranks to transition from strong HLA-binder (SB, netMHCpan %Rank < 0.5) to HLA non-binders (NB, netMHCpan %Rank > 2.0) were considered as leading to ‘Loss of binding’. Mutations causing predicted binding affinities to transition from NB to SB were considered as leading to ‘Gain of binding’.

### Selection of clinically validated CD8+ T-Cell epitopes

A list of validated CD8+ T Cell epitopes presented by both HLA-A and -B molecules were downloaded from https://www.mckayspcb.com/SARS2TcellEpitopes/ (as of January 2021). This database, developed by Dr. Matthew R. McKay and his team, contains compiled and catalogued validated T-cell epitope-HLA pairs from 13 studies aimed at identifying immunogenic SARSCOV-2 T-cell epitopes.

### In vitro HLA-peptide binding assays

Peptide binding to class I HLA molecules was quantitatively measured using classical competition assays based on the inhibition of binding of a high affinity radiolabeled peptide to purified HLA molecules, as detailed elsewhere (Sidney et al., 2013). Briefly, HLA molecules were purified from lysates of EBV transformed homozygous cell lines by affinity chromatography by repeated passage over Protein A Sepharose beads conjugated with the W6/32 (anti-HLA-A, -B, -C) antibody, following separation from HLA-B and -C molecules by pre-passage over a B1.23.2 (antiHLA B, C) column. Protein purity, concentration, and the effectiveness of depletion steps was monitored by SDS-PAGE and BCA assay. Peptide affinity for respective class I molecules was determined by incubating 0.1-1 nM of radiolabeled peptide at room temperature with 1 µM to 1 nM of purified HLA in the presence of a cocktail of protease inhibitors and 1 µM B2microglobulin. Following a two-day incubation, HLA bound radioactivity was determined by capturing MHC/peptide complexes on W6/32 antibody coated Lumitrac 600 plates (Greiner Bioone, Frickenhausen, Germany). Bound cpm was measured using the TopCount (Packard Instrument Co., Meriden, CT) microscintillation counter. The concentration of peptide yielding 50% inhibition of the binding of the radiolabeled peptide was calculated. Under the conditions utilized, where [label]<[MHC] and IC50 ≥ [MHC], the measured IC50 values are reasonable approximations of the true Kd values. Each competitor peptide was tested at six different concentrations covering a 100,000-fold dose range, and in three or more independent experiments. As a positive control for inhibition, the unlabeled version of the radiolabeled probe was also tested in each experiment.

### SANTA-SIM simulations

We simulated SARS-CoV-2 genomes with SANTA-SIM, using the consensus sequence WuhanHu-1 as input sequence available at https://www.ncbi.nlm.nih.gov/nuccore/MN908947.3. Each simulation was run with a population size of 10,000 individual viral sequences evolving for 1000 generations, and analyses were conducted on random samples of 1,000 viral sequences. Following Huddelston et.al. (Huddleston et al., 2020) who used SANTA-SIM to simulate influenza A/H3N2 that has a yearly substitution rate approximately twice as high as SARS-CoV-2 [∼48,824 substitutions/year (https://nextstrain.org/flu/seasonal/h3n2/ha/2y?l=clock) vs. ∼24.5 substitution/year (https://nextstrain.org/ncov/global?l=clock)], we chose 400 generations/year, with the mutation rate per position per generation set to 2.04E-6 (yearly substitution rate/(generations in one year * genome size)). The transition bias was set to 3.0 for baseline simulations. To evaluate the impact of specific substitution biases, additional simulations were conducted using a substitution matrix with scores set to 1.0 of transversions, 3.0 for transitions, and biases ranging from 4.0 to 20.0 for the targeted substitution. We generated 10 replicates for all simulated scenarios, except for C-to-U where we made 100 replicates to better assess statistical significance.

### Determination of amino acid mutational patterns

Mutational biases were identified by calculating the overall change in amino acid composition caused by the mutational landscape of SARS-CoV-2 for each individual amino acid, referred in the main text as ‘global residue substitution output’ (GRSO). For this analysis, all mutations found globally in at least 4 GISAID entries were analysed together. Preferential introduction or removal of amino acids was determined by comparing the overall amino acid composition in reference residues vs mutated residues throughout the mutation pool, resulting in a percentile difference in amino acid composition. As such, for amino acid *X*, the % difference was calculated according to the following formula:

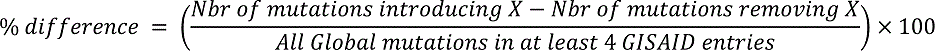

This analysis took into consideration the number of unique mutations. Therefore, to consider mutational biases in the context of mutation frequencies, the analysis described above was conducted separately for mutations occurring in a single GISAID entry (expected to be enriched for errors); 2-10 GISAID entries; 11-99 GISAID entries; and 100 or more GISAID entries. As a negative control, the SANTA SIM algorithm was used to simulate the neutral evolution of 1000 SARS-CoV-2 genomes (baseline simulations, N = 10 replicates). This control was used to calculate the statistical significance of the observed biases, by way of a One-Sample T-Test.

### Prediction of mutation impacts on peptide presentation in the context of HLA supertypes

Reference/mutated peptide pairs for which the differential predicted binding affinities led to transitions from strong HLA binder (SB) to non-HLA binder (NB) [(SB) NetMHCpan %rank < 0.5 to (NB) NetMHCpan %rank >2] or from NB to SB, were identified, catalogued and analyzed as described above. Binding affinities were predicted for representative HLA types from several major HLA supertypes (A1, A2, A3, A24, B7, B8, B27, B44), as defined by Sydney *et al*. We then categorized all reference/mutated peptide pairs on the basis of their 1) mutation type (amino acid *X* à amino acid *Y*) and 2) the position of the mutation in the peptide sequence. Finally, we quantified the number of reference/mutated peptide pairs and the associated average fold change in predicted binding affinity for each category. P-values were generated for each category by performing a two-tailed independent T-Test between the fold changes in binding affinity associated with mutation type *A* at position *X*, and all fold changes in binding affinity associated with position *X*.

### Assessing the contribution of nucleic acid mutation types to the global amino acid mutational patterns

To assess the contribution of various nucleic acid mutation types to the observed amino acid mutational patterns, we first determined the respective contributions of each nucleic acid mutation type to the global mutation landscape. We then selected the five most abundant mutation types [CàU (41%), GàU (18%), AàG, GàA, UàC (9.7-11.6%)] and assessed their individual impacts on amino acid mutational patterns using the simulation algorithm SANTA SIM as follows: For each mutation type, we simulated the evolution of 1000 SARS-CoV-2 genomes over 1000 generations (N = 10 replicates) with varying degrees of biases (the coefficient used to determine the extent of the biases was exploratively set to ‘x4’, ‘x8’, ‘x15’, and ‘x20’) (Figure S6A). Because the input coefficient does not have a linear relationship with the abundance of the mutation type observed in the simulation output, we used the simulations with all four parameter values (x4, x8, x15, x20) in order to identify the simulation parameter that most closely reflected observations in real-life SARS-CoV-2 data. The coefficient for the ratio of *X* à *Y* nucleic acid mutation type to all other mutation types was generated using the following formula:

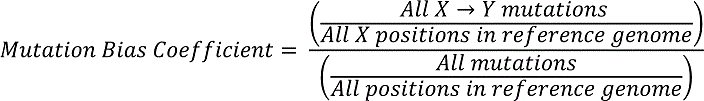

Finally, all amino acid mutations were identified for the output of each simulation, as described above. To determine statistical significances, simulated mutational biases (at the amino acid level) were compared to a neutral evolution as a negative control (N = 10 replicates) by way of twotailed independent T-Test.

### Statistical analysis

A Two-tailed One-Sample T-Test was used to assess the statistical significance of the observed mutational biases against the neutral simulations (N = 10 replicates). A Two-tailed Independent T-Test assuming different variances was used to assess the statistical significances of 1) the simulated biased SARS-CoV-2 evolution, 2) the gain/loss plots in the context of supertypes, and 3) the statistical significance associated with the average fold change in %rank associated with each position-specific amino acid mutation type in the supertype analysis.

## SUPPLEMENTARY MATERIALS

### SUPPLEMENTARY FIGURE LEGENDS

**Figure S1.**
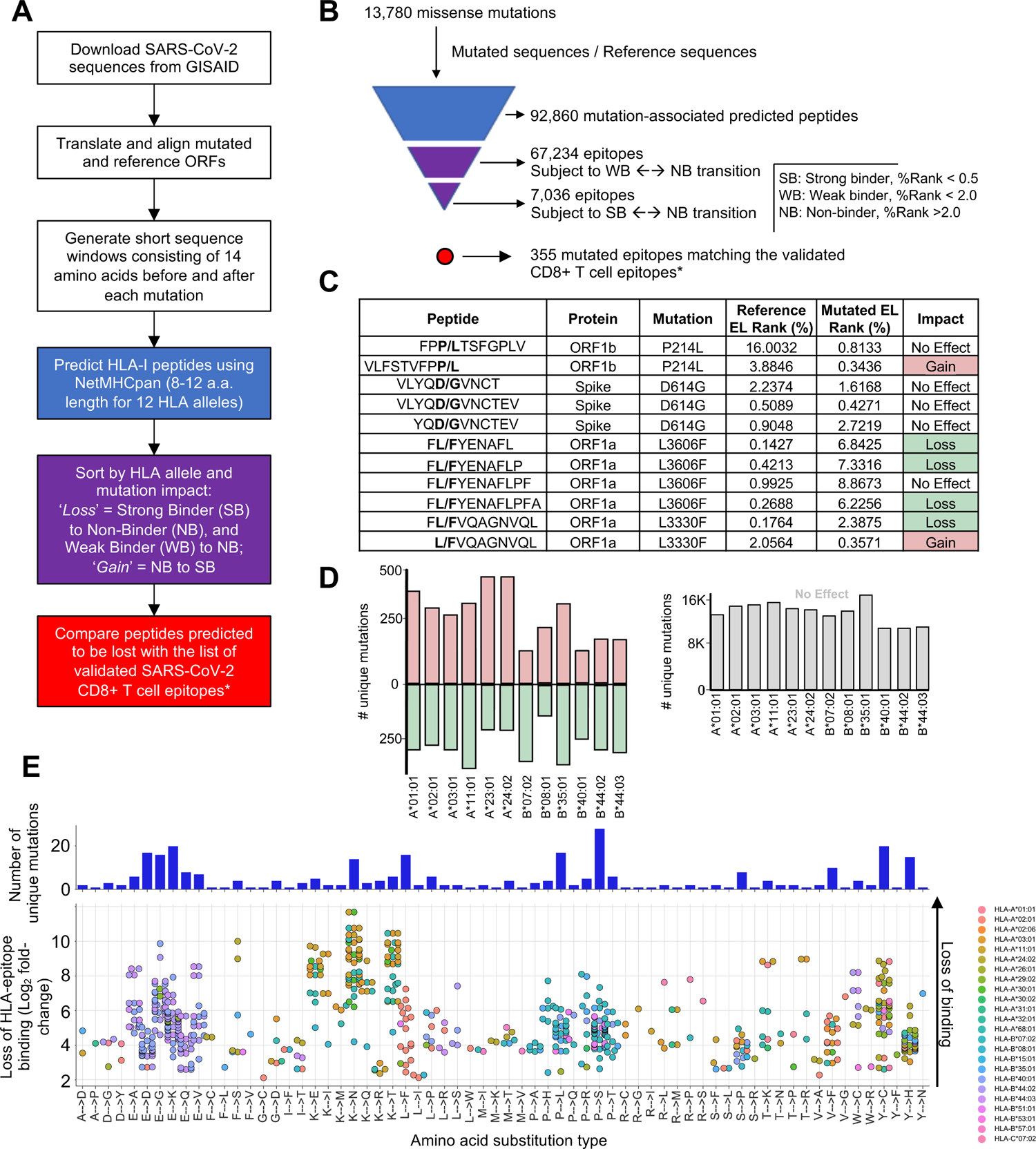
Impact of SARS-CoV-2 mutations on CD8+ T cell epitopes, Related to. Figure 1 **and 4**. (**A**) Bioinformatic pipeline for the prediction of SARS-CoV-2 mutated class I peptides associated to 12 common HLA alleles. (**B**) Pyramidal graph showing the number of i) missense mutations in SARS-CoV-2 genomes, ii) predicted class I mutated peptides, iii) predicted class I peptides subject to Weak Binder (WB) to Non-Binder (NB) and Strong Binder (SB) to NB transition (epitope loss category), and iv) predicted class I mutated peptides matching reference CD8+ T cell epitopes that have been experimentally validated. (**C**) Representative examples of predicted class I mutated peptides and the impact of the identified amino acid mutation (bold) on peptide binding to a given HLA-I allele. Reference and mutated EL (eluted ligand) Rank (%) generated by NetMHCpan 4.1 EL is indicated for individual predictions. Gain = NB to SB (pale red); Loss = SB to NB (pale green). (**D**) Left panel: number of unique mutations leading to ‘Gain’ or ‘Loss’ of class I peptides for the indicated HLA-I alleles. Right panel: number of unique mutations showing no effect on peptide binding for the indicated HLA-I alleles. (**E**) Validated SARS-CoV-2 CD8+ T cell epitopes (McKay Database) subjected to mutation events detected in more than 4 individuals (GISAID) and predicted lead to a strong loss of HLA-epitope binding. Top: number of unique missense mutations corresponding to the indicated amino acid substitution type. Bottom: Predicted loss of HLA-epitope binding (NetMHCpan4.1 %Rank) corresponding to the indicated residue substitution type from the list of validated CD8+ T cell epitopes in the McKay Database. Each dot represents an epitope pair (mutated / reference). Color indicates HLA type affected by the mutations.

**Figure S2.**
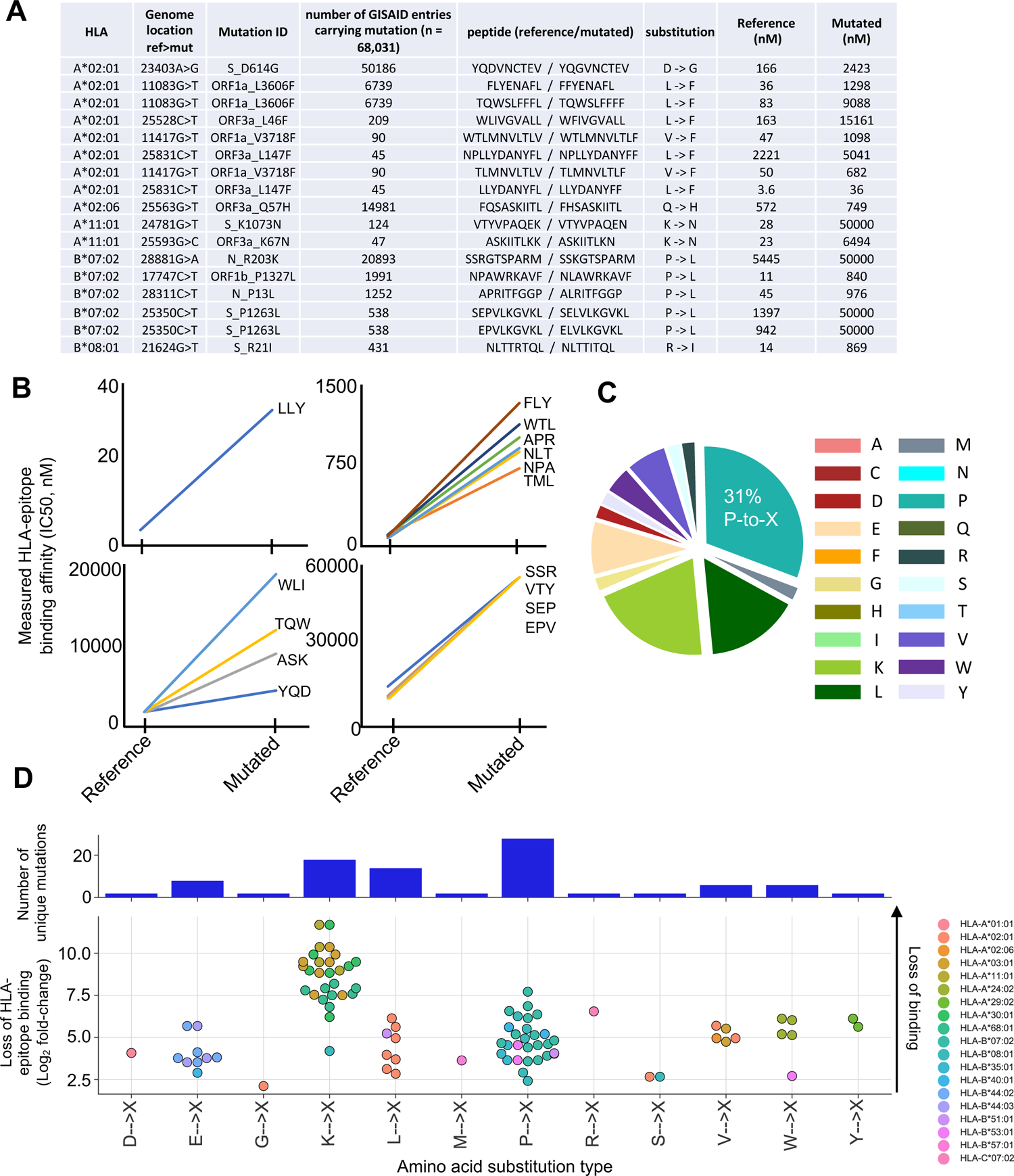
HLA peptide binding measurements and mutational biases in SARS-CoV-2 mutated epitopes, Related to. Figure 1. (**A**) HLA binding assay was performed to determine the in vitro binding affinity (nM) of representative SARS-CoV-2 peptides for specific HLA class I alleles. Peptides were selected based on 1) frequency of mutations, 2) presentation by common HLA class I alleles, and 3) the mutated form was predicted to lose binding to its corresponding HLA. (**B**) Plots showing raw values for the binding affinities (nM) of the reference vs mutated peptides in (A). The first three amino acid residues of the reference peptides with fold change > 2.5 are shown. (**C**) Pie chart showing the proportion of X-to-Y substitution types from the list of validated CD8+ T cell epitopes in https://www.mckayspcb.com/SARS2TcellEpitopes/ (as of January 2021). (**D**) Predicted loss of HLA-epitope binding clustered by substitution type from the list of validated CD8+ T cell epitopes in the McKay database. Each dot represents an epitope pair (mutated / reference; NetMHCpan 4.1 %rank ratio).

**Figure S3.**
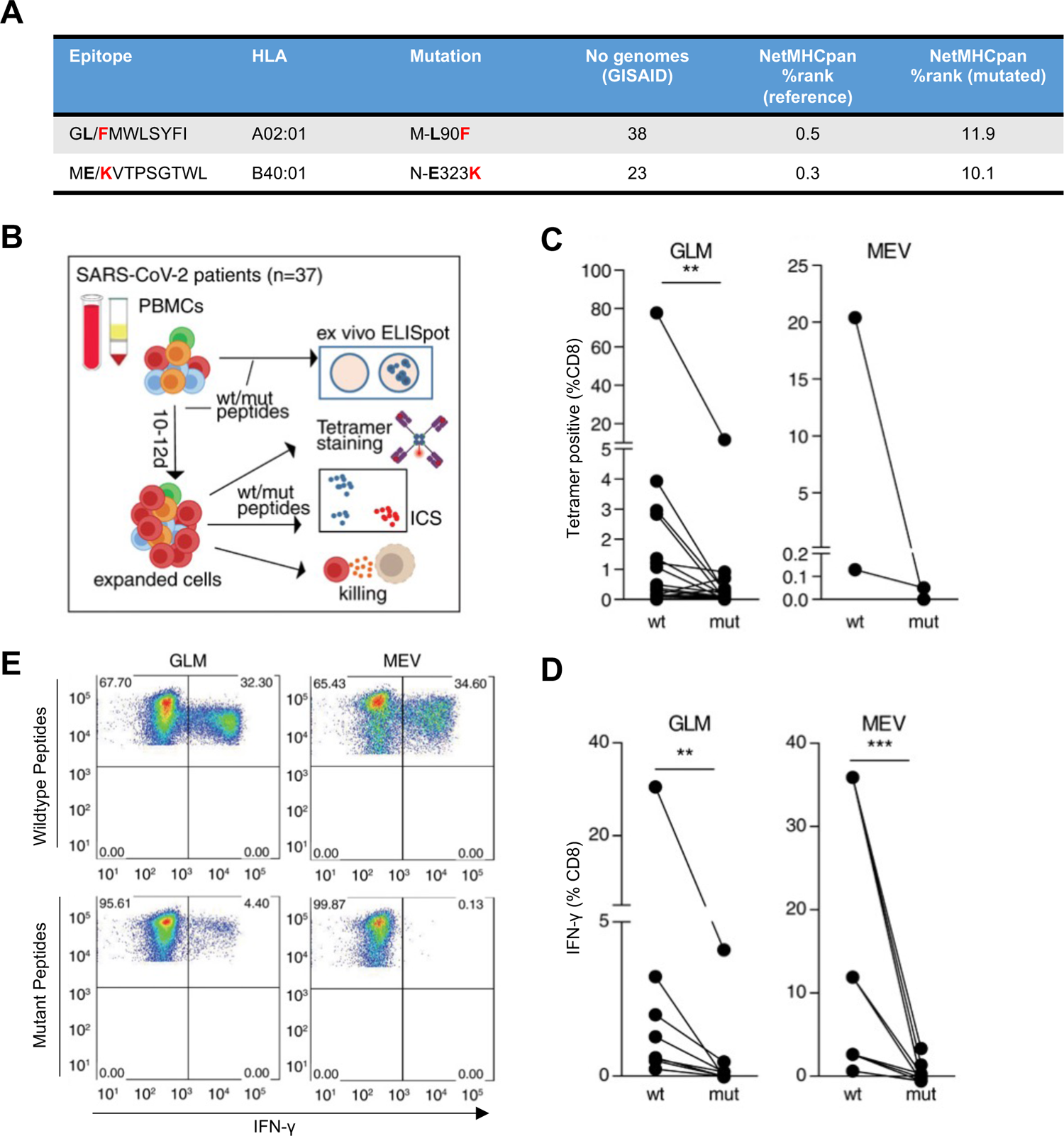
Identification of two SARS-CoV-2 mutated epitopes in this study that were previously associated with decreased CD8+ T cell responses, Related to. Figure 1. (**A**) The mutated epitopes GFMWLSYFI (A*02) and MKVTPSGTWL (B*40) were detected in 38 and 23 genomes/individuals in this study (GISAID) and their T cell immunogenicity was thoroughly investigated in Agerer et al. (**B-E from Agerer et al., copyright 2021, with permission from AAAS)** (**B**) Experimental overview. (**C**) T cells expanded with mutant peptides do not give rise to wild type peptide-specific CD8+ T cell. PBMCs were isolated from HLA-A*02:01 or HLA-B*40:01 positive SARS-CoV-2 patients, stimulated with wild type or mutant peptides and stained with tetramers containing the wild type peptide. (**D**) Impact of mutations on CD8+ T cell response. PBMCs expanded with wild type or mutant peptides as indicated, were analyzed for IFN-γ-production via ICS after restimulation with wild type or mutant peptide. (**E**) Representative FACS plots for (D).

**Figure S4.**
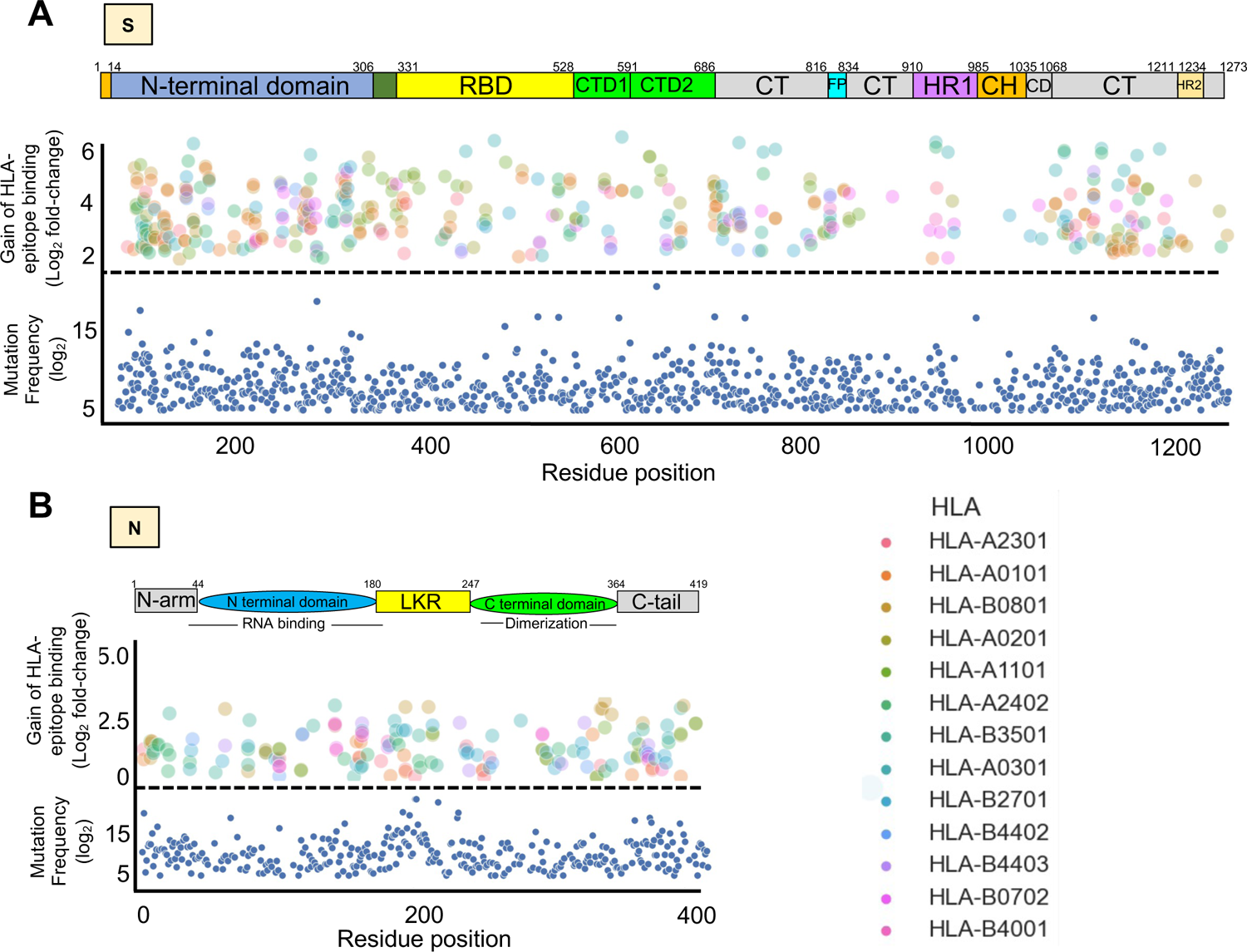
Impact of mutations on gain of peptide binding to various HLA class I molecules across the immunodominant S and N antigens, Related to. Figure 1. (**A, B**) Lower panel: blue dots showing all mutations that occurred in at least 4 SARS-CoV-2 genomes (GISAID). Upper panel: dots showing predicted peptides subjected to a strong gain of binding (see also Figure S1C,D) to one of 12 highly common HLA types queried (color coded) due to a mutation.

**Figure S5.**
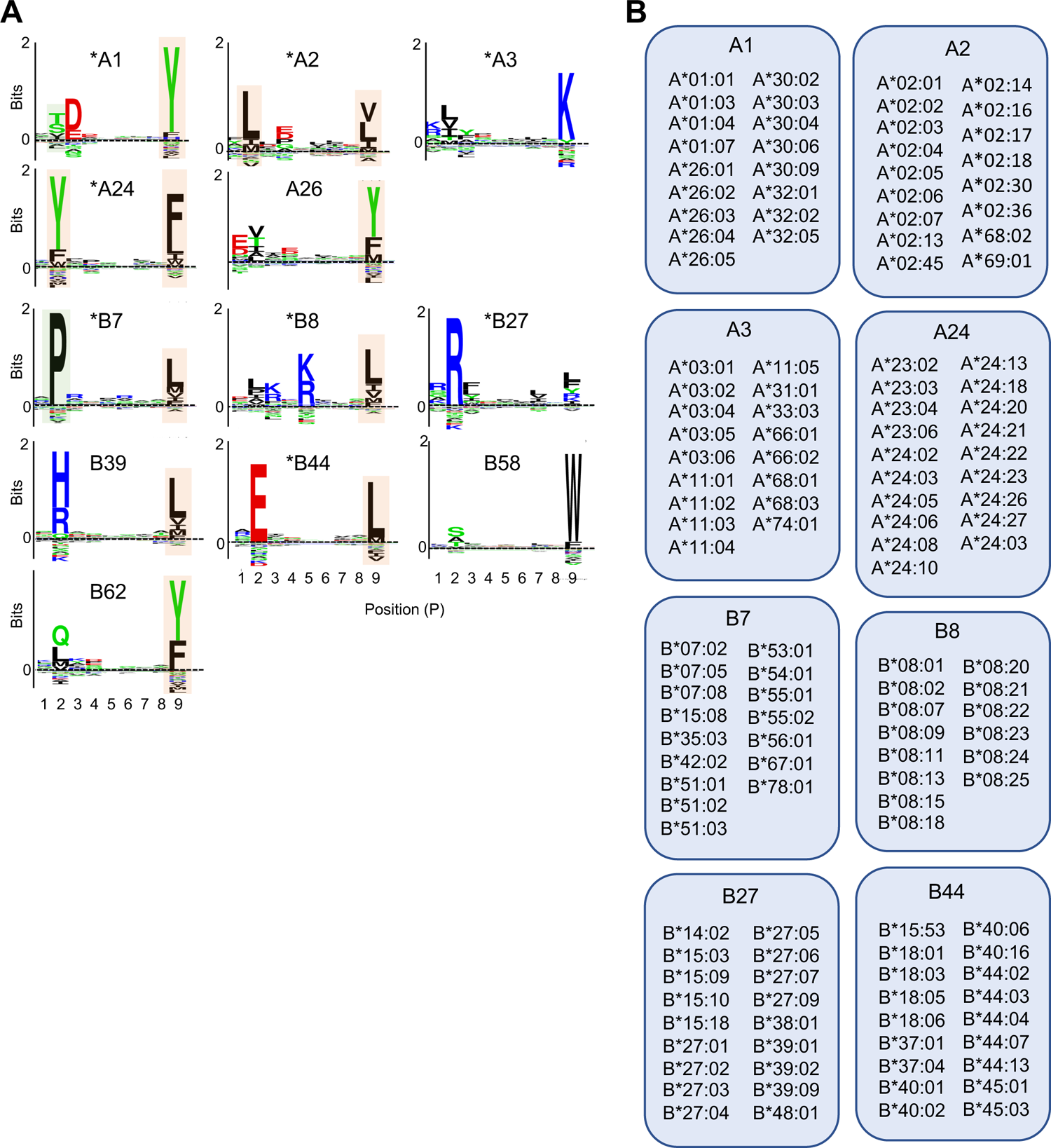
HLA class I supertypes, Related to. Figure 4. (**A**) Epitope binding motifs for several HLA class I supertypes. Anchor residues are located at P2 and P9. Pale orange and green squares cover amino acid residues that are preferentially introduced (F, I, L, Y) and removed (A, P, T) in SARS-CoV-2 proteomes, respectively. Representative supertypes used in this study are shown by an asterisk. Epitope binding motifs were extracted from NetMHCpan Motif Viewer (http://www.cbs.dtu.dk/services/NetMHCpan/logos_ps.php). (**B**) Table showing the selected alleles per supertype that were used in this study to generate the ‘Gain/Loss plots’.

**Figure S6.**
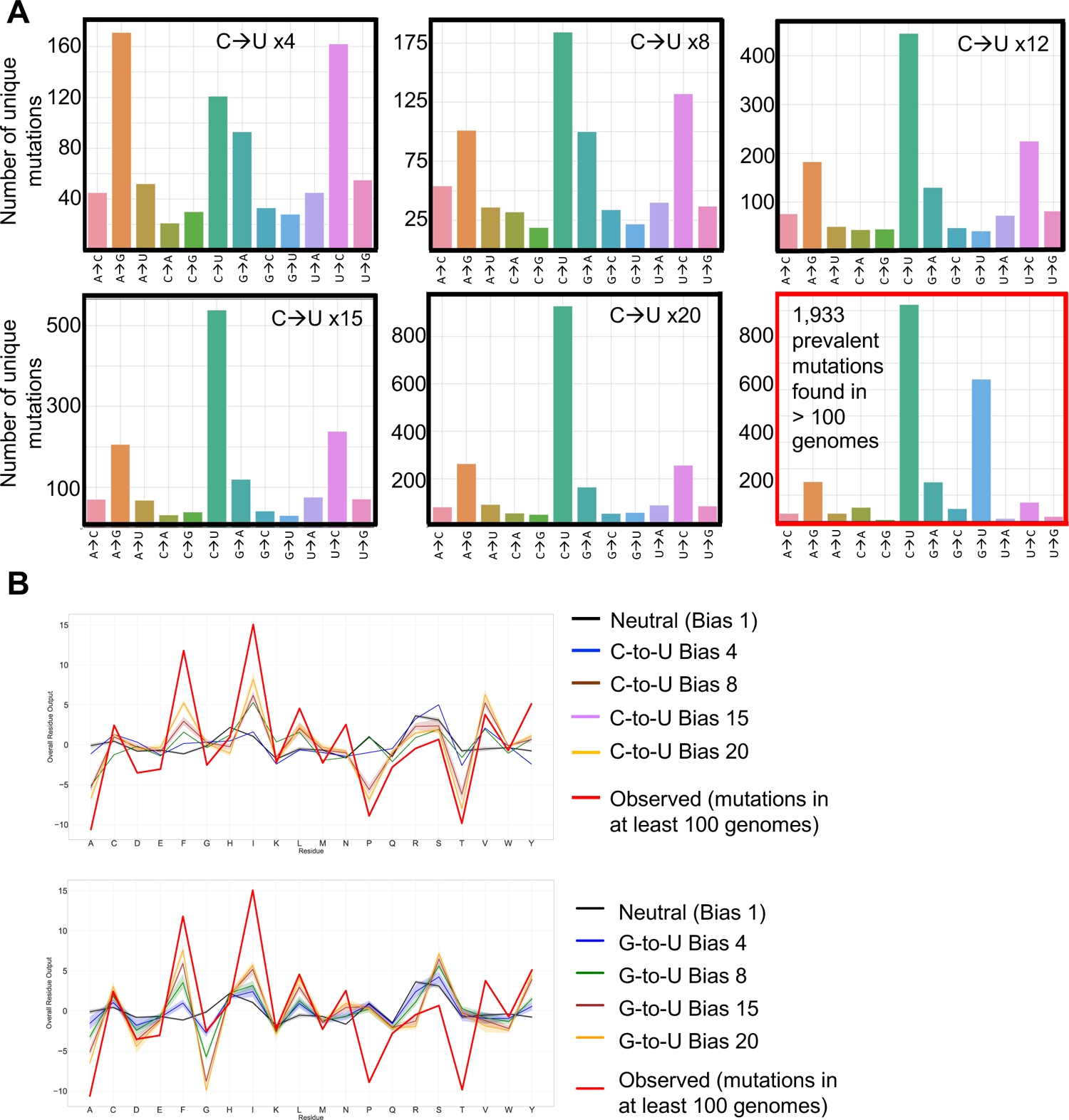
Comparison of mutation biases between real-life/observed and simulated data, Related to. Figure 5. (**A**) Histograms showing the number of unique mutations identified for each mutation type (A-to-C, A-to-G, etc.) after simulating the evolution of SARS-CoV-2 genomes through the introduction of different C-to-U bias values (x4 to x20) using the SANTA-SIM software. Simulated (black squares) and real-life/observed prevalent mutations found in more than 100 genomes (red square) at the nucleotide level are shown. (**B**) Comparison of global amino acid mutational patterns generated from simulated versus real-life/observed SARS-COV-2 genomes. Various extents of C-to-U (top) and G-to-U (bottom) biases were introduced to perform the simulation and to generate the graphs.

### SUPPLEMENTARY TABLE LEGENDS

**Table S1. SARS-CoV-2 mutations identified from 330,246 GISAID entries (December 31^st^ 2020), Related to Figure 1**. SARS-CoV-2 mutations at the nucleic and amino acid level are indicated. Number of genomes carrying mutation show the frequency of individual mutations among all SARS-CoV-2 variants.

**Table S2.** SARS-CoV-2 prevalent mutations identified from 330,246 GISAID entries (December 31^st^ 2020) and detected in at least 100 individuals, Related to Figure 1.

**Table S3.** Documented SARS-CoV-2 CD8+ T cell epitopes and their matching mutated forms identified in this study, Related to **Figure 1**.

**Table S4. List of documented SARS-CoV-2 CD8+ T cell epitopes.** Epitopes were downloaded from https://www.mckayspcb.com/SARS2TcellEpitopes/ (as of January 2021). This database has effectively catalogued all SARS-CoV-2 CD8+ epitopes validated by 18 separate studies.

